# Structure-based Generation of a Secondary Nucleation Inhibitor in α-Synuclein Aggregation Using a Conditional Diffusion Model

**DOI:** 10.1101/2025.08.16.670694

**Authors:** Hao Zhang, Robert I. Horne, Z. Faidon Brotzakis, Charles Harris, Pietro Liò, Michele Vendruscolo

## Abstract

The process of *α*-synuclein aggregation results in the formation of amyloid fibrils, which accumulate in the brain of patients affected by Parkinson’s disease. Among possible therapeutic strategies to cure this disease, one approach is based on the development of compounds capable of inhibiting *α*-synuclein aggregation. An effective inhibition could be achieved by blocking the nucleation sites on the surface of the amyloid fibrils that are responsible for their autocatalytic proliferation. Here, we report a strategy based on deep learning to achieve this goal, which uses an E(3)-equivariant conditional diffusion model. By using this approach, we designed and tested experimentally candidate small molecules. We found that one of these small molecules acts as a potent inhibitor of secondary nucleation in *α*-synuclein aggregation. These results provide evidence that generative diffusion models offer effective tools for drug design.

## Introduction

Parkinson’s disease is the most common neurodegenerative movement disorder. ^1–5^ Because the disease is characterised by the presence of aberrant deposits formed by *α*-synuclein, ^6,7^ many efforts have been focused on understanding the links between the aggregation of this protein and pathological processes in Parkinson’s disease.^8,9^ Consequently, *α*-synuclein has emerged as a natural target for drug discovery. ^10–12^

The aggregation of *α*-synuclein takes place through a complex kinetic process, which involves a network of coupled microscopic steps. ^13,14^ Small oligomeric assemblies form initially from monomeric precursors through primary nucleation, in a process often catalysed by lipid membranes. ^14,15^ The oligomers, which are initially disordered, can convert into ordered forms, ^16^ which can grow into long fibrils through monomer-dependent elongation. ^13,14^ The presence of these fibrils can then catalyse the formation of new oligomeric assemblies, in an autocatalytic process responsible for the rapid proliferation of *α*-synuclein deposits. ^13,14,17^ This process is known as secondary nucleation because it depends on the presence of already formed amyloid fibrils, and it is typically much faster than primary nucleation.^18^

Recent advances in chemical kinetics approaches have allowed the identification of small molecules and molecular chaperones that are able to inhibit *α*-synuclein aggregation. ^19–24^ It has thus been possible to specifically inhibit the surface-catalysed secondary nucleation step, which is responsible for the autocatalytic proliferation of *α*-synuclein fibrils, by binding competitively with *α*-synuclein monomers along specific sites on the surface of *α*-synuclein fibrils. ^20,23,25,26^ Considering that oligomers are generated primarily by surface-catalysed secondary nucleation,^27^ the discovery of compounds targeting this mechanism offers promising opportunities for drug discovery. ^20,23,26,28^

Structure-based drug design, which can be used to target the catalytic site for secondary nucleation, is typically carried out by using virtual screening methods, ^29–31^ most commonly by molecular docking.^32^ This approach has been recently implemented using ultra-large chemical libraries of billions of compounds. ^33–35^

Here, we explore a different route, by building on the observation that small molecule generation by deep learning is an efficient way to explore the chemical space, and it can be used as an alternative to virtual screening ^36–41^ in the initial steps of drug discovery. The problem of generating small molecules for drug design can be split up into three components: (i) generation of a small molecule, (ii) fitting the small molecule into a binding pocket, and (iii) endowment of desired properties (e.g. potency, selectivity, synthesizability and drug-likeliness) into the small molecule. The achievement of these objectives can be assessed using benchmarks. ^42–44^

An important aspect of generative models for small molecules is the molecular representation that they adopt. This choice is important to perform predictions by maximizing the information translated into the latent representation downstream of the neural network architecture. Broadly speaking, representations can be one-dimensional (1D, typically strings), two-dimensional (2D, typically graphs), or three-dimensional (3D, typically coordinates). Initial efforts focused on 1D representations, such as SMILES (Simplified Molecular Input Line Entry System), ^45^ which represent small molecules as strings of characters, were based on already available neural network architectures for language processing. ^36^ However, it is still challenging to incorporate valence and synthesizability rules into this type of approach. ^46^ Moreover, small changes in SMILES may cause a large changes in the small molecules. To overcome these challenges, more recent work has focused on 2D representations, by representing small molecules as graphs, with nodes corresponding to atoms, and edges to bonds. This strategy captures in a natural manner the connectivity between atoms, but requires the development of specialised neural network architectures. The rapid development of graph neural networks has greatly helped this task. ^47,48^ To capture not only the connectivity but also the different possible conformations of small molecules, recent work has focused on 3D representations, where atoms are represented through their coordinates.^49,50^

Another relevant aspect of generative models is in the type of neural networks used to build small molecules. Currently used approaches include variational autoencoders (VAE), ^36,51,52^ generative adversarial networks (GAN),^53–55^ normalizing flows, ^56–59^ re-current neural networks (RNN), ^60–62^ graph neural networks (GNNs),^48,63^ and generative pretrained transformers. ^64–67^

Diffusion models have been recently applied to address this problem in a novel manner. ^68–72^ This technique comprises two steps - a forward noising process to transform a data distribution into pure noise and backward process to remove the noise and reconstruct the starting distribution. Diffusion models can then be used to generate new samples starting from pure noise. This approach has recently also been implemented using neural operators. ^73^ Diffusion models have been recently used for molecular generation. ^71,72,74–78^ Here we adapt a recent approach, ^79^ which offers an implementation of the inductive bias that the shape of the building pocket should match that of the ligand.

Our results show that our strategy can be used to generate a potent inhibitor of *α*-synuclein aggregation, which, according to a series of in vitro aggregation assays, is of comparable potency to compounds currently in clinical trials based on a similar mechanism of action.

## Results

### Choice of the binding pocket

In this work, we targeted a binding pocket on the structure of an amyloid fibril of *α*-synuclein recently reported as a catalytic site for secondary nucleation^26^ (Figure 1A). This binding pocket, which comprises residues His50-Lys58 and Thr72-Val77, was identified by comparing two structures determined using cryo-electron microscopy, one obtained in vitro using a recombinant protein (PDB ID: 6cu7) and the other obtained from brain-derived samples (PDB ID: 8a9l) (Figure 1B,C). This binding pocket was originally determined using Fpocket, ^80^ which identifies potential binding pockets based on volume criteria (Figure 1A), and CamSol, which provides a solubility score, ^81^ and then validated experimentally by showing that small molecules binding to it inhibit secondary nucleation.^26^ Moreover, *α*-synuclein secondary nucleation has been reported as significant only below pH 5.8, when His50 is protonated, also supporting the choice of the surface binding pocket. ^27^

**Figure 1.**
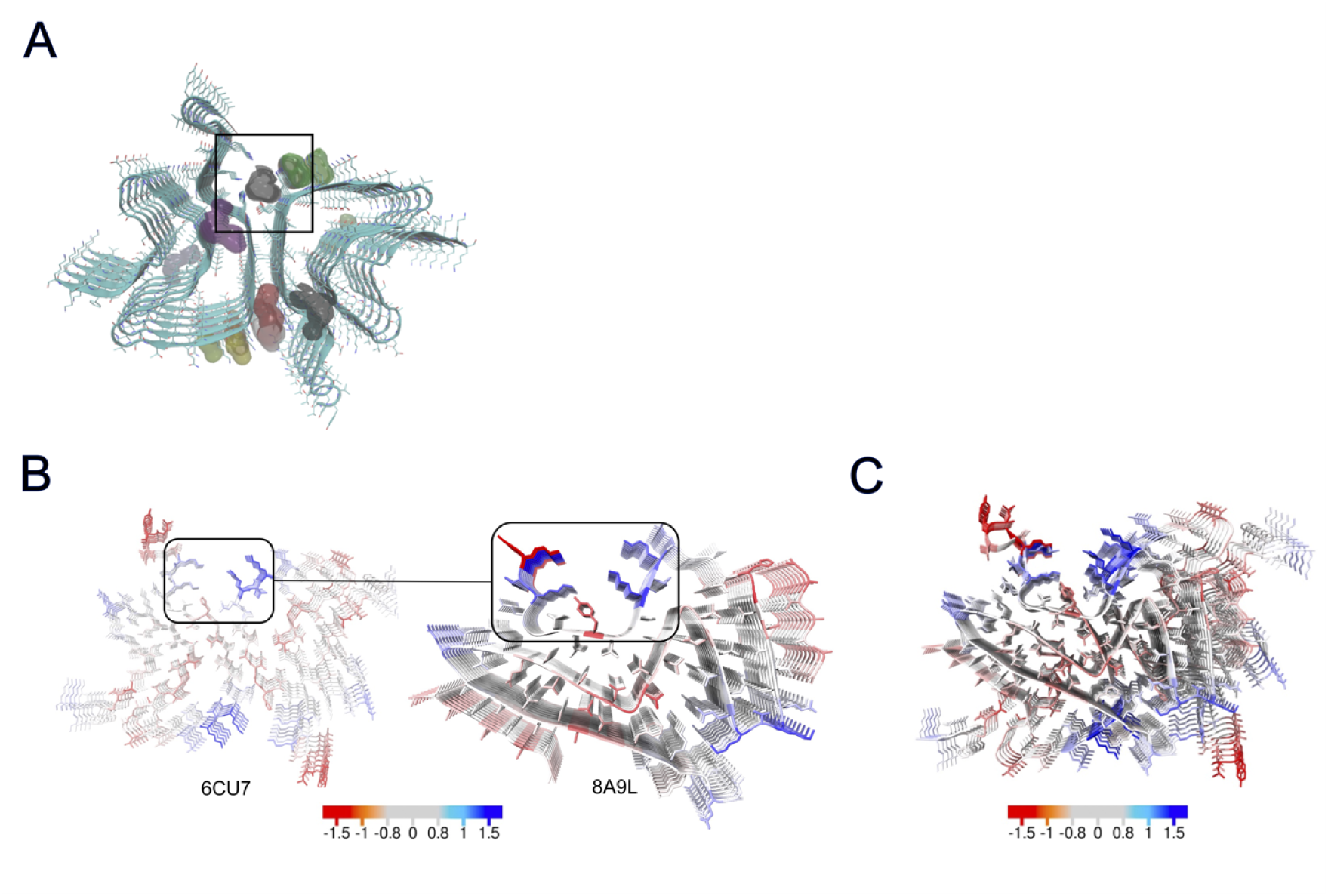
Illustration of the binding pocket on an α-synuclein fibril targeted in this study. (A) Binding pocket prediction based on Fpocket^80^ on the cryo-EM structures of a recombinant amyloid fibril of *α*-synuclein (6cu7). This binding pocket was recently validated experimentally by showing that small molecules binding to it inhibit secondary nucleation.^26^ (B) Comparison of the structures of the recombinant amyloid fibril described in panel A with a brain derived one (8a9l). The homologous binding site is indicated by a square box. (C) Structural overlap of amyloid fibril structures in panel B, with the binding sites aligned. The structures are coloured according to the CamSol residue solubility score.^81^ Adapted from.^26^

### Candidate compounds

We adopted a multistep strategy to design and test candidate compounds. These steps are flexible, and they can be readily changed for adapting them to other targets. Firstly, we adapted a recently proposed conditional model from,^79^ which in turn was based on a previous model. ^74^ Details for the model are provided in the Supplementary Information. We used this model to generate small molecules conditioned to bind the target binding pocket.

We then filtered out molecules predicted to bind to the target binding pocket with low affinity using the Vina score from Autodock Vina. ^82^ From the molecules generated by the diffusion model, we selected the 12 molecules with Vina score lower than -7 kcal/mol, as lower scores indicate better binding affinity.

Since the synthesis of small molecules is costly and time consuming, and not guaranteed to be achievable when the small molecules themselves are obtained through generative modelling, it would be beneficial if the candidate compounds could be available for purchase via vendors. We thus searched for small molecules in the ZINC database that are similar to the 12 selected small molecules. The similarity was calculated based on topological fingerprint and Tanimoto similarity, where hydrogen atoms are not explicitly considered. We only considered molecules with a molecular weight lower than 500 Da, thus selecting 300 molecules from the ZINC database with the highest similarity to the 12 selected small molecules. By filtering out those molecules with Vina score lower than -7 kcal/mol, we obtained 129 candidate compounds (Table S1). Among these, we ordered 7 for experimental testing, which had the best Vina scores and were readily available for purchase.

### Experimental validation of the candidate compounds

We tested experimentally a set of seven small molecules generated through the procedure explained above. The structures and experimental anti-aggregation metrics for these molecules are shown in Figure 2 and Table S2.

**Figure 2.**
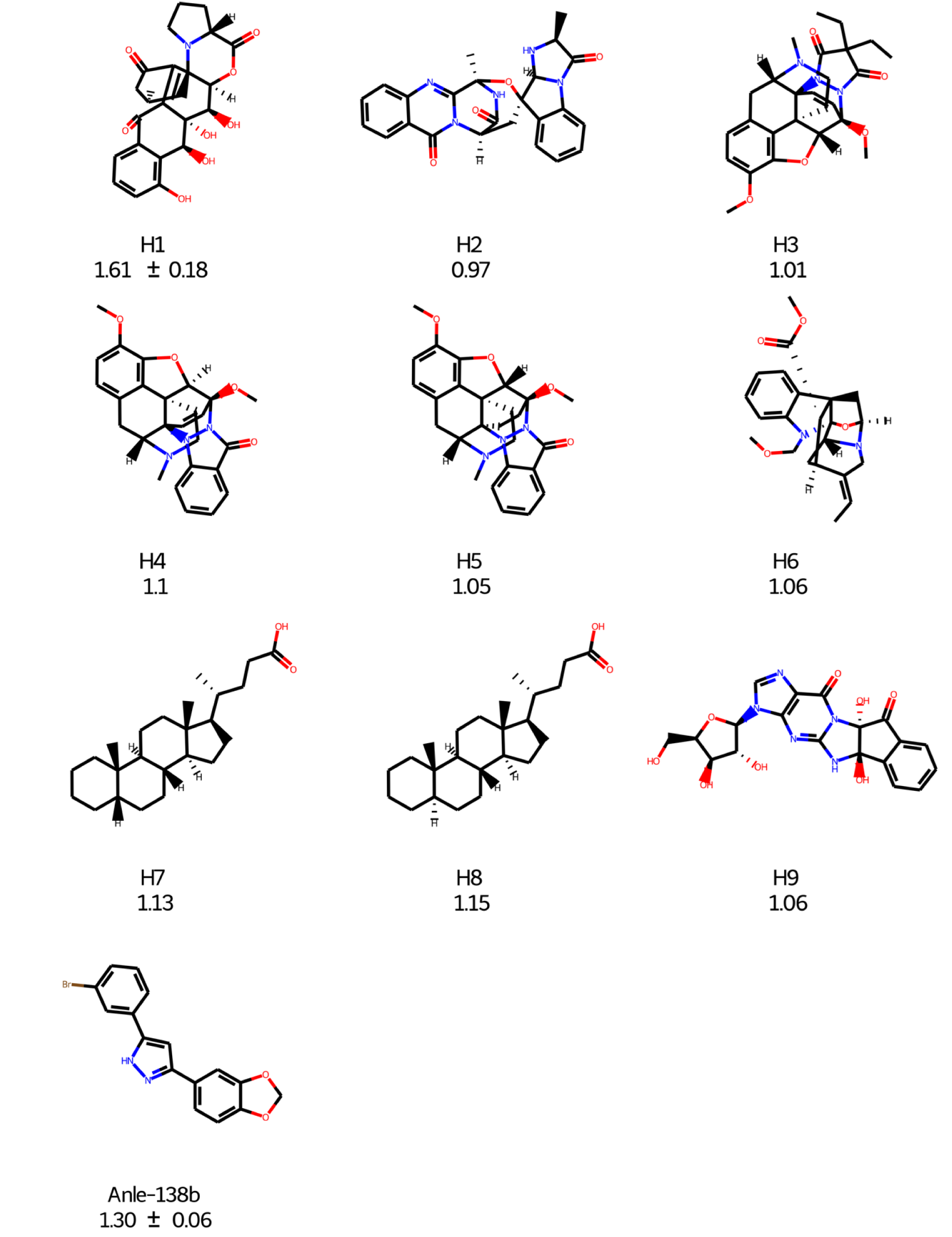
Structures of the 7 compounds tested experimentally. The fold-increase in the half time of aggregation is indicated below each structure, alongside the compound label. For comparison, we also show Anle-138b.

We employed a chemical kinetics assay designed to identify key compounds inhibiting the surface-catalysed secondary nucleation phase within the aggregation process of *α*-synuclein ^26,83^ (Figure 3). The negative control, containing 1% DMSO, is shown in dark purple, and the positive control, containing Anle-138b,^84^ in blue. Anle-138b is currently in clinical trials for inhibition of *α*-synuclein aggregation and exhibits efficacy in the assay used here.

**Figure 3.**
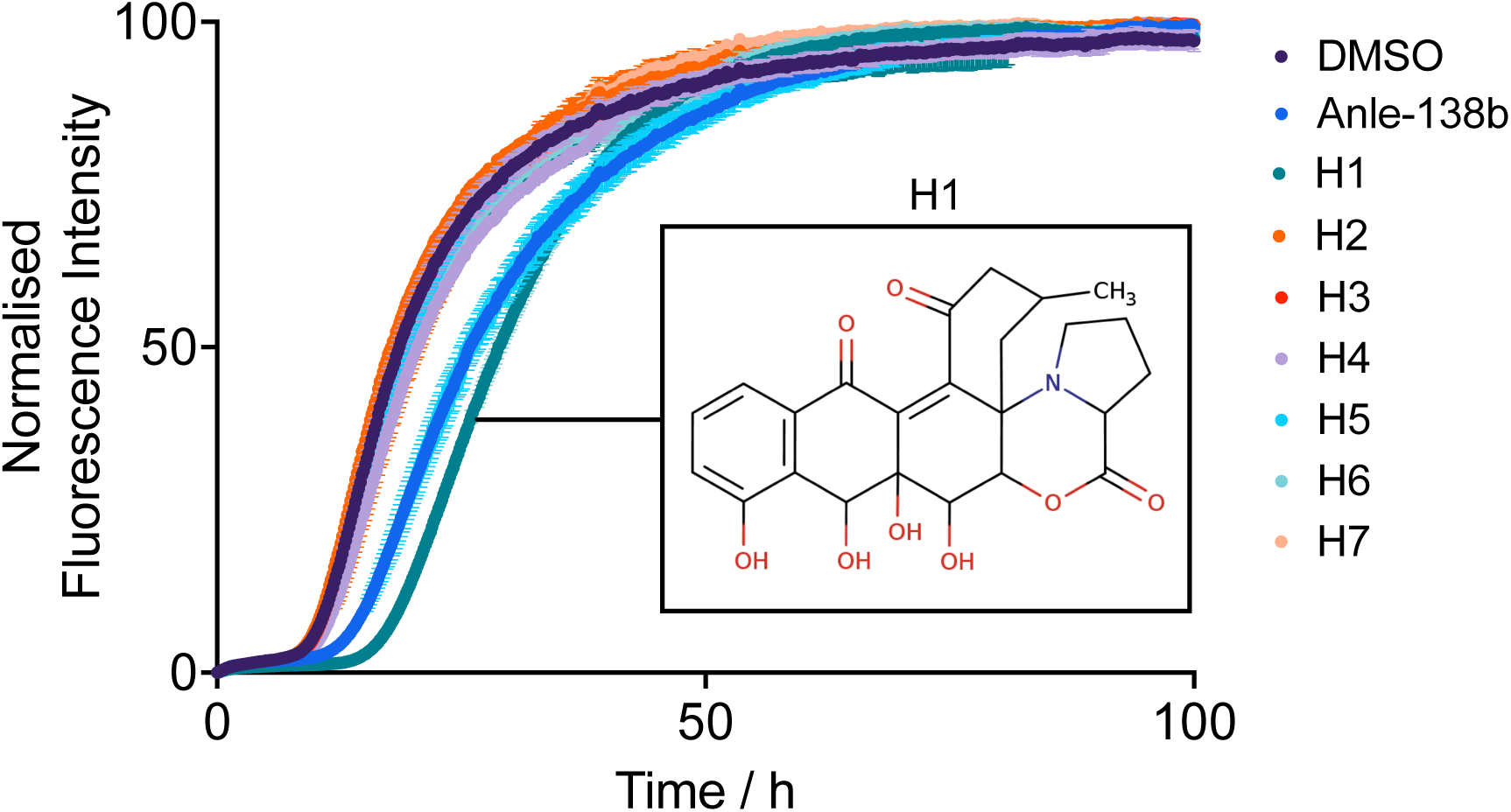
In vitro aggregation assay of the 7 predicted inhibitors selected for testing. Kinetic traces of a 10 *µ*M solution of *α*-synuclein with 25 nM seeds at pH 4.8, 37 °C in the presence of the compounds (25 *µ*M) or 1% DMSO (n=2 replicates, central measure=mean, error=SD). The negative control (1% DMSO) is shown in purple, and the positive control (Anle-138b in 1% DMSO) is shown in blue. The structure of the most potent inhibitor (H1) is also shown. In this experiment, H1 exhibited higher potency than Anle-138b, a compound currently in clinical trials as inhibitor of *α*-synuclein aggregation^84^

To run the aggregation assay under conditions in which secondary nucleation is the dominant aggregation mechanism, we introduced a small quantity of preformed fibrils into a monomeric mixture at the start of the reaction. ^26,27^ One of the seven small molecules tested, H1, was found to have significant potency, with normalised half time of 1.6 ± 0.2 h (25 *µ*M H1, mean and SD from 2 independent experiments).

This half time is normalised to that of the negative control, giving a value of 1 for the control. For comparison, the normalised half time of Anle-138b was 1.3 ± 0.1 h (25 *µ*M Anle-138b, mean and SD from 2 independent experiments).

To analyse in more detail the inhibitory effect of H1, we carried out further aggregation experiments using increasing concentrations of H1, which confirmed that this small molecule exhibits a dose-dependent effect on *α*-synuclein aggregation (Figure 4).

**Figure 4.**
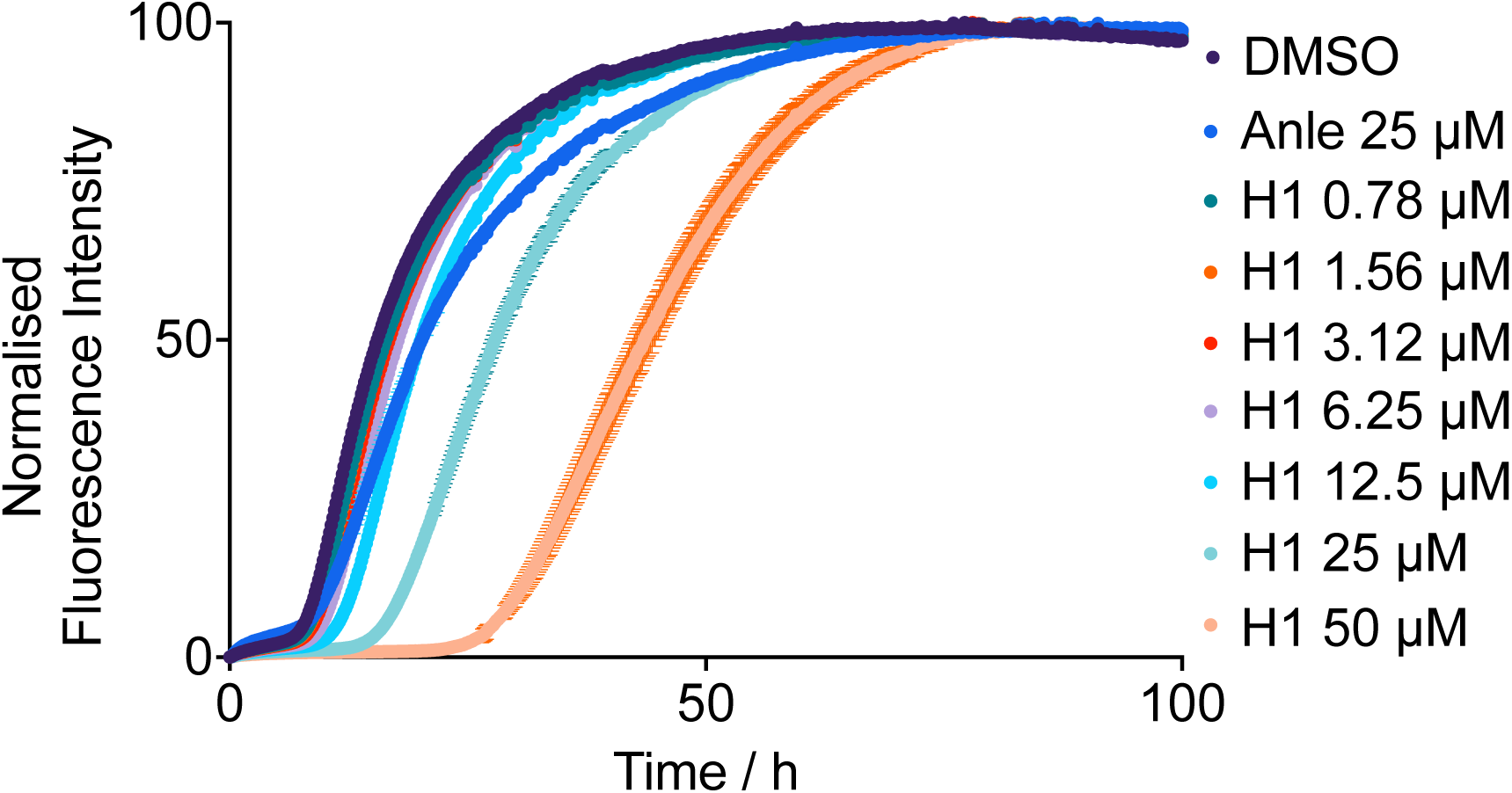
Dose dependence of the effect of compound H1 on the in vitro aggregation assay of α-synuclein. Kinetic traces of a 10 *µ*M solution of *α*-synuclein with 25 nM seeds at pH 4.8, 37 °C in the presence of molecule or 1% DMSO (n=2 replicates, central measure=mean, error=SD). The negative control is shown in purple, and the positive control (Anle-138b, 25 *µ*M) in blue. The structure of the most potent inhibitor (H1) is also shown.

## Conclusions

We presented a deep learning method of designing small molecule inhibitors of *α*-synuclein aggregation based on a conditional diffision model. We provided evidence that this strategy can generate compounds with a specific mechanism of action, namely targeting catalytic sites for secondary nucleation on amyloid fibrils to prevent the proliferation of aggregates. These results illustrate the potential of deep learning methods based on diffusion models to facilitate the process of drug design.

## Methods

### Diffusion model

In this work, we used a diffusion method recently reported, ^79^ which in turn was based on a previous model. ^74^ A summary of method is reported in the Supplementary Information.

### Compounds and chemicals

Compounds were purchased from MolPort (Riga, Latvia) and prepared in DMSO to a stock of 5 mM. All chemicals used were purchased at the highest purity available.

### Recombinant ***α***-synuclein expression

Recombinant *α*-synuclein was purified based on previously described methods. ^13,14,85^ The plasmid pT7-7 encoding human *α*-synuclein was transformed into BL21 (DE3) competent cells. Following transformation, the competent cells were grown in 6L 2xYT media in the presence of ampicillin (100 *µ*g/mL). Cells were induced with IPTG, grown overnight at 28 °C and then harvested by centrifugation in a Beckman Avanti JXN-26 centrifuge with a JLA-8.1000 rotor at 6240 rcf (Beckman Coulter, Fullerton, CA). The cell pellet was resuspended in 10 mM Tris, pH 8.0, 1 mM EDTA, 1 mM PMSF and lysed by sonication. The cell suspension was boiled for 20 min at 85 °C and centrifuged at 39,000 relative centrifugal force (rcf) with a JA-25.5 rotor (Beckman Coulter). Streptomycin sulfate was added to the supernatant to a final concentration of 10 mg/mL and the mixture was stirred for 15 min at 4 °C. After centrifugation at 39,000 rcf, the supernatant was taken with an addition of 0.36 g/mL ammonium sulfate. The solution was stirred for 30 min at 4 °C and centrifuged again at 39,000 rcf. The pellet was resuspended in 25 mM Tris, pH 7.7, and the suspension was dialysed overnight in the same buffer. Ion-exchange chromatography was then performed using a Q Sepharose HP column of buffer A (25 mM Tris, pH 7.7) and buffer B (25 mM Tris, pH 7.7, 1.5 M NaCl). The fractions containing *α*-synuclein were loaded onto a HiLoad 26/600 Superdex 75 pg Size Exclusion Chromatography column, and the protein (≈ 60 ml @ 200 *µ*M) was eluted into the required buffer. The protein concentration was determined spectrophotometrically using the extinction coefficient *ε*280 = 5600 M-1 cm-1.

### ***α***-synuclein seed fibril preparation

*α*-synuclein fibril seeds were produced as described previously. ^13,85^ Samples of *α*-synuclein (700 *µ*M) were incubated in 20 mM phosphate buffer (pH 6.5) for 72 h at 40 °C and stirred at 1,500 rpm with a Teflon bar on an RCT Basic Heat Plate (IKA, Staufen, Germany). Fibrils were then diluted to 200 *µ*M, aliquoted and flash frozen in liquid N2, and finally stored at -80 °C. For the use of kinetic experiments, the 200 *µ*M fibril stock was thawed, and sonicated for 15 s using a tip sonicator (Bandelin, Sonopuls HD 2070, Berlin, Germany), using 10% maximum power and a 50% cycle.

### Measurement of ***α***-synuclein aggregation kinetics

*α*-synuclein was injected into a Superdex 75 10/300 GL column (GE Healthcare) at a flow rate of 0.5 mL/min and eluted in 20 mM sodium phosphate buffer (pH 4.8) supplemented with 1 mM EDTA. The obtained monomer was diluted in buffer to a desired concentration and supplemented with 50 *µ*M ThT and preformed *α*-synuclein fibril seeds. The molecules (or DMSO alone) were then added at the desired concentration to a final DMSO concentration of 1% (v/v). Samples were prepared in low-binding Eppendorf tubes, and then pipetted into a 96-well half-area, black/clear flat bottom polystyrene NBS microplate (Corning 3881), 150 *µ*L per well. The assay was then initiated by placing the microplate at 37 °C under quiescent conditions in a plate reader (FLUOstar Omega, BMG Labtech, Aylesbury, UK). The ThT fluorescence was measured through the bottom of the plate with a 440 nm excitation filter and a 480 nm emission filter. After centrifugation at 2350 rcf to remove aggregates the monomer concentration was measured via the Pierce™ BCA Protein Assay Kit according to the manufacturer’s protocol.

## Notes

### Competing Interest Statement

The authors have declared no competing interest.

