## Supplementary Information for "Structure-based Generation of a Secondary Nucleation Inhibitor in α-Synuclein Aggregation Using a Conditional Diffusion Model"

### Diffusion Models

We introduce diffusion models for the common case of discrete unconditional diffusion models with Gaussian noise.[1–3] We use $q$ to indicate probabilities for fixed distributions and $p_\theta$ for distributions changing with a parameter $\theta$.

A data point and its conditions are represented as a vector $\mathbf{x}$. Gaussian noise is added to the data points to obtain $\mathbf{z}_t$ in $t$ steps, and $t \in \{t \mid 1 \leq t \leq T, t \in \mathbb{Z}\}$. This process is referred to as diffusion. $\alpha_t \in \mathbb{R}^+$ and $\sigma_t \in \mathbb{R}^+$ control this diffusion process so that

$$q\left(\mathbf{z}_t \mid \mathbf{x}\right) = \mathcal{N}\left(\mathbf{z} \mid \alpha_t \mathbf{x}, \sigma_t^2 \mathbf{I}\right). \tag{1}$$

Typically, $\alpha_t = \sqrt{1-\sigma_t^2}$, as in variance preserving processes. Based on this diffusion process, one can obtain

$$q\left(\mathbf{z}_{t2} \mid \mathbf{z}_{t1}\right) = \mathcal{N}\left(\mathbf{z}_{t2} \mid \frac{\alpha_{t2}}{\alpha_{t1}}\mathbf{z}_{t1}, \left(\sigma_{t2}^2 - \frac{\alpha_{t2}^2}{\alpha_{t1}^2}\sigma_{t1}^2\right)\mathbf{I}\right), \tag{2}$$

in which $t1 < t2$.

The diffusion process is then paired with a denoising process. Since the distribution $q\left(\mathbf{z}_{t1} \mid \mathbf{z}_{t2}\right)$ is not tractable, one considers instead the distribution

$$q\left(\mathbf{z}_{t1} \mid \mathbf{z}_{t2}, \mathbf{x}\right) = \mathcal{N}\left(\mathbf{z}_{t1} \mid \frac{\alpha_{t2}\sigma_{t1}^2}{\alpha_{t1}\sigma_{t2}^2}\mathbf{z}_{t2} + \left(\alpha_{t1} - \frac{\alpha_{t2}^2\sigma_{t1}^2}{\alpha_{t1}\sigma_{t2}^2}\right)\mathbf{x}, \left(\sigma_{t1}^2 - \frac{\alpha_{t2}^2\sigma_{t1}^4}{\alpha_{t1}^2\sigma_{t2}^2}\right)\mathbf{I}\right) \tag{3}$$

When we use the diffusion model to do sampling, we do not have $\mathbf{x}$, but we can use a neural network to estimate $\mathbf{x}$ as $\hat{\mathbf{x}}$. $\hat{\mathbf{x}}$ can be obtained from a neural network.[2] It is also possible to estimate the noise added to $\mathbf{z}_{t2}$ as $\hat{\boldsymbol{\epsilon}}$ for easier optimization.[3] In this work, we also estimate $\hat{\boldsymbol{\epsilon}}$. The relationship between $\hat{\boldsymbol{\epsilon}}$ and $\hat{\mathbf{x}}$ is

$$\hat{\mathbf{x}}_\theta = \frac{1}{\alpha_{t2}}\mathbf{z}_{t2} - \frac{\sigma_{t2}}{\alpha_{t2}}\hat{\boldsymbol{\epsilon}}_\theta. \tag{4}$$

Then, we can get

$$p_\theta\left(\mathbf{z}_{t1} \mid \mathbf{z}_{t2}\right) = \mathcal{N}\left(\mathbf{z}_{t1} \mid \frac{\alpha_{t2}\sigma_{t1}^2}{\alpha_{t1}\sigma_{t2}^2}\mathbf{z}_{t2} + \left(\alpha_{t1} - \frac{\alpha_{t2}^2\sigma_{t1}^2}{\alpha_{t1}\sigma_{t2}^2}\right)\hat{\mathbf{x}}_\theta, \left(\sigma_{t1}^2 - \frac{\alpha_{t2}^2\sigma_{t1}^4}{\alpha_{t1}^2\sigma_{t2}^2}\right)\mathbf{I}\right). \tag{5}$$

In choosing the loss function for model training, we aim to reduce the negative log-likelihood of $p_\theta\left(\mathbf{x}\right)$. However, it is not tractable directly, so we use the evidence lower bound (ELBO) instead, getting

$$-\log p_\theta\left(\mathbf{x}\right) \leq L_{ELBO} = \mathbb{E}_q\left(L_x + \Sigma_{t=2}^T L_t + L_T\right). \tag{6}$$

When the diffusion process is fixed, i.e. it has no learnable parameter, also

$$L_T = D_{\mathrm{KL}}\left(q\left(\mathbf{z}_T \mid \mathbf{x}\right) \| \mathcal{N}\right) \tag{7}$$

is fixed, and we can ignore this term during training. The other two terms are

$$L_x = -\log p_\theta\left(\mathbf{x} \mid \mathbf{z}_1\right) \tag{8}$$

$$L_t = D_{\mathrm{KL}}\left(q\left(\mathbf{z}_{t-1} \mid \mathbf{z}_t, \mathbf{x}\right) \| p_\theta\left(\mathbf{z}_{t-1} \mid \mathbf{z}_t\right)\right). \tag{9}$$

$L_x$ can be viewed as a reconstruction term, and it is related to data representation. $L_t$ is the main focus point. Typically, according to,[3] these two terms can be further simplified, and we can optimize a simpler loss:

$$\mathbb{E}\|\hat{\boldsymbol{\epsilon}}_\theta - \boldsymbol{\epsilon}\|^2. \tag{10}$$

The pseudocodes for the unconditional model is provided as Algorithm 1 and Algorithm 2. The change from an unconditional model to a conditional model is straightforward. Condition $\mathbf{c}$ is added to the input of the neural network, and the format is changed from $\hat{\boldsymbol{\epsilon}}_\theta\left(\mathbf{z}_t, t\right)$ to $\hat{\boldsymbol{\epsilon}}_\theta\left(\mathbf{z}_t, t, \mathbf{c}\right)$.

**Algorithm 1** Training for the unconditional diffusion model.

```
1: repeat
2:     x ~ q(x)
3:     t ~ U({t | 1 ≤ t ≤ T, t ∈ Z})
4:     ε ~ N(0, I)
5:     z_t = α_t x + σ_t ε
6:     Gradient descent on ||ε̂_θ(z_t, t) − ε||^2
7: until converged
```

### Atom-Only Model

In an atom-only model no chemical bonds are included, and all the chemical bonds in the results are added during post-process by Open Babel.[4]

**Algorithm 2** Sampling for the unconditional diffusion model.

1: $\mathbf{z}_T \sim \mathcal{N}(0, \mathbf{I})$
2: **for** $t = T, T-1, \cdots, 3, 2$ **do**
3: $\quad \boldsymbol{\epsilon} \sim \mathcal{N}(0, \mathbf{I})$
4: $\quad \hat{\mathbf{x}}_\theta = \frac{1}{\alpha_t}\mathbf{z}_t - \frac{\sigma_t}{\alpha_t}\hat{\boldsymbol{\epsilon}}_\theta(\mathbf{z}_t, t)$
5: $\quad \mathbf{z}_{t-1} = \frac{\alpha_t \sigma_{t-1}^2}{\alpha_{t-1}\sigma_t^2}\mathbf{z}_t + \left(\alpha_{t-1} - \frac{\alpha_t^2 \sigma_{t-1}^2}{\alpha_{t-1}\sigma_t^2}\right)\hat{\mathbf{x}}_\theta + \sqrt{\sigma_{t-1}^2 - \frac{\alpha_t^2 \sigma_{t-1}^4}{\alpha_{t-1}^2 \sigma_t^2}}\boldsymbol{\epsilon}$
6: $\mathbf{x} \sim p_\theta(\mathbf{x} \mid \mathbf{z}_1)$

#### Data Representation

In this work, we used a full atom model, which means that the protein binding pockets are represented in the same way as the small molecule ligands. They are represented in a point cloud format with atom coordinates $\mathbf{C} \in \mathbb{R}^{n\times 3}$ and atom type features $\mathbf{A} \in \{0, 1\}^{n\times t_a}$, in which $n$ is the number of atoms and $t_a$ is the number of atom types. We use $\mathbf{X} = [\mathbf{C}_0, \mathbf{A}_0]$ to represent the raw data and $\mathbf{Z}_t = [\mathbf{C}_t, \mathbf{A}_t], t > 1$ for data with noise. $(P)$ and $(L)$ in the superscript refer to pocket and ligand respectively.

#### Diffusion and Denoising

The diffusion process and denoising process are equivariant to permutation and rigid transformation. The detail of the diffusion process and denoising process is shown in Algorithm 3 and Algorithm 4. During these processes, $\mathbf{C}$ and $\mathbf{A}$ are normalized by scaling to allow the neural network learning more easily, and coordinates are centred by removing the mean of coordinates to make the process equivariant, i.e. the output follows the same transformation rules of the input, for instance under rigid transformation.

To estimate the noise, we used the following neural network

$$\hat{\boldsymbol{\epsilon}}_\theta\left(\mathbf{Z}_t^{(L)}, t, \mathbf{X}^{(P)}\right), \tag{11}$$

in which $\mathbf{X}^{(P)}$ is the condition for the diffusion model. The neural network used here is an Equivariant Neural Network (EGNN) from.[5] The features for the ligands and the pockets

**Algorithm 3** Training for the atom-only model.

**repeat**

  $\mathbf{X} \sim q(\mathbf{X})$

  $t \sim U(\{t \mid 1 \leq t \leq T, t \in \mathbb{Z}\})$

  $\boldsymbol{\epsilon} \sim \mathcal{N}(0, \mathbf{I})$

  Normalizing $\mathbf{C}$ and $\mathbf{A}$

  $\mathbf{C}_0 = \mathbf{C}_0 - \text{mean}\left(\mathbf{C}_0^{(L)}\right)$

  $\mathbf{Z}_t^{(L)} = \alpha_t \mathbf{X}^{(L)} + \sigma_t \boldsymbol{\epsilon}$

  $\mathbf{C}_0^{(P)} = \mathbf{C}_0^{(P)} - \text{mean}\left(\mathbf{C}_t^{(L)}\right)$

  $\mathbf{C}_t^{(L)} = \mathbf{C}_t^{(L)} - \text{mean}\left(\mathbf{C}_t^{(L)}\right)$

  Gradient descent on $\|\hat{\boldsymbol{\epsilon}}_\theta\left(\mathbf{Z}_t^{(L)}, t, \mathbf{X}^{(P)}\right) - \boldsymbol{\epsilon}\|^2$

**until** converged

**Algorithm 4** Sampling for the atom-only model.

**Require:** $\mathbf{X}^{(P)}$

Normalizing $\mathbf{C}_0^{(P)}$ and $\mathbf{A}_0^{(P)}$

$\mathbf{C}_T^{(L)} \sim \mathcal{N}\left(\text{mean}\left(\mathbf{C}_0^{(P)}\right), \mathbf{I}\right)$

$\mathbf{C}_T = \mathbf{C}_T - \text{mean}\left(\mathbf{C}_T^{(L)}\right)$

$\mathbf{A}_T^{(L)} \sim \mathcal{N}(0, \mathbf{I})$

**for** $t = T, T-1, \cdots, 3, 2$ **do**

  $\boldsymbol{\epsilon} \sim \mathcal{N}(0, \mathbf{I})$

  $\hat{\mathbf{X}}_\theta^{(L)} = \frac{1}{\alpha_t}\mathbf{Z}_t^{(L)} - \frac{\sigma_t}{\alpha_t}\hat{\boldsymbol{\epsilon}}_\theta\left(\mathbf{Z}_t^{(L)}, t, \mathbf{X}^{(P)}\right)$

  $\mathbf{Z}_{t-1}^{(L)} = \frac{\alpha_t \sigma_{t-1}^2}{\alpha_{t-1}\sigma_t^2}\mathbf{Z}_t^{(L)} + \left(\alpha_{t-1} - \frac{\alpha_t^2 \sigma_{t-1}^2}{\alpha_{t-1}\sigma_t^2}\right)\hat{\mathbf{X}}_\theta^{(L)} + \sqrt{\sigma_{t-1}^2 - \frac{\alpha_t^2 \sigma_{t-1}^4}{\alpha_{t-1}^2 \sigma_t^2}}\boldsymbol{\epsilon}$

  $\mathbf{C}_0^{(P)} = \mathbf{C}_0^{(P)} - \text{mean}\left(\mathbf{C}_{t-1}^{(L)}\right)$

  $\mathbf{C}_{t-1}^{(L)} = \mathbf{C}_{t-1}^{(L)} - \text{mean}\left(\mathbf{C}_{t-1}^{(L)}\right)$

$\mathbf{X}^{(L)} \sim p_\theta\left(\mathbf{X}^{(L)} \mid \mathbf{Z}_1^{(L)}, \mathbf{X}^{(P)}\right)$

Unnormalizing $\mathbf{C}_0^{(L)}$ and $\mathbf{A}_0^{(L)}$

Onehot encode $\mathbf{A}_0^{(L)}$

Undo centering

are embedded, and all features and time are concatenated together.

$$\begin{bmatrix} \vdots \\ \mathbf{a}_i \\ \vdots \\ \mathbf{a}_j \\ \vdots \end{bmatrix} = \begin{bmatrix} \text{embed}\left(\mathbf{A}_t^{(L)}\right), \mathbf{t}^{(L)} \\ \text{embed}\left(\mathbf{A}_0^{(P)}\right), \mathbf{t}^{(P)} \end{bmatrix}, \mathbf{t}^{(L)} = \begin{bmatrix} t \\ \vdots \\ t \end{bmatrix}, \mathbf{t}^{(P)} = \begin{bmatrix} t \\ \vdots \\ t \end{bmatrix} \tag{12}$$

All coordinates are also concatenated together.

$$\begin{bmatrix} \vdots \\ \mathbf{c}_i \\ \vdots \\ \mathbf{c}_j \\ \vdots \end{bmatrix} = \begin{bmatrix} \mathbf{C}_t^{(L)} \\ \mathbf{C}_0^{(P)} \end{bmatrix} \tag{13}$$

The updating rules for each layer are

$$\mathbf{m}_{ij} = \phi_m\left(\mathbf{a}_i^l, \mathbf{a}_j^l, d_{ij}^2\right) \tag{14}$$

$$e_{ij} = \phi_{\text{att}}\left(\mathbf{m}_{ij}\right) \tag{15}$$

$$\mathbf{a}_i^{l+1} = \phi_f\left(\mathbf{a}_i^l, \sum e_{ij}\mathbf{m}_{ij}\right) \tag{16}$$

$$\mathbf{c}_i^{l+1} = \mathbf{c}_i^l + \sum \frac{\mathbf{c}_i^l - \mathbf{c}_j^l}{d_{ij}+1}\phi_c\left(\mathbf{a}_i^l, \mathbf{a}_j^l, d_{ij}^2\right) + \frac{\left(\mathbf{c}_i^l - \bar{\mathbf{c}}^l\right) \times \left(\mathbf{c}_j^l - \bar{\mathbf{c}}^l\right)}{\left\|\left(\mathbf{c}_i^l - \bar{\mathbf{c}}^l\right) \times \left(\mathbf{c}_j^l - \bar{\mathbf{c}}^l\right)\right\| + 1}\phi_c'\left(\mathbf{a}_i^l, \mathbf{a}_j^l, d_{ij}^2\right), \tag{17}$$

in which $\phi_m$, $\phi_{\text{att}}$, $\phi_f$, $\phi_c$, and $\phi_c'$ are MLP, $d_{ij}$ is the distance. If there is any edge feature, it can be passed to MLPs along with distance. $\sum$ in the equation apply to edges shorter than the distance cutoff. There are several ($L$) such layers, and $\mathbf{A}$ is embedded before and after all these layers (another twice in total). At the output stage, $\mathbf{X}^{\text{out}} = \mathbf{X}^L - \mathbf{X}^0$, and $\mathbf{A}$ is embedded again after removing the time part. All embeddings are done with MLPs or

SLPs. Only the noise estimated for ligands is kept.

Table S1 **List of the top 129 candidate compounds from the ZINC database.** We provide the Vina scores (in kcal/mol), the molecular weight (in Da), the number of atoms, and the ZINC identifier.

| smiles | vina score | weight | num atom | ZINC id |
|---|---|---|---|---|
| CN1C(=O)[C@H]2C[C@]3(O[C@H]1c1nc4ccccc4c(=O)n12)c1ccccc1N1C(=O)C2(CC2)N[C@@H]13 | -59.97 | 455.1593541520006 | 55 | 1772591296 |
| CN1C(=O)[C@H]2C[C@@]3(O[C@H]1c1nc4ccccc4c(=O)n12)c1ccccc1N1C(=O)C2(CC2)N[C@@H]13 | -59.67 | 455.1593541520006 | 55 | 1772591294 |
| CN1C(=O)[C@H]2C[C@]3(O[C@H]1c1nc4ccccc4c(=O)n12)c1ccccc1N1C(=O)C2(CC2)N[C@H]13 | -59.43 | 455.1593541520006 | 55 | 1772591295 |
| CN1C(=O)[C@H]2C[C@@]3(O[C@H]1c1nc4ccccc4c(=O)n12)c1ccccc1N1C(=O)C2(CC2)N[C@H]13 | -59.31 | 455.1593541520006 | 55 | 1772591297 |
| C[C@H]1C(=O)N2c3ccccc3[C@@]3(O)C[C@@H]4C(=O)N[C@](C)(c5nc6ccccc6c(=O)n54)N1[C@@H]23 | -17.66 | 443.1593541520007 | 54 | 103216961 |
| C[C@H]1C(=O)N2c3ccccc3[C@@]3(O)C[C@@H]4C(=O)N[C@@](C)(c5nc6ccccc6c(=O)n54)N1[C@@H]23 | -15.49 | 443.1593541520007 | 54 | 95786121 |
| C[C@@H]1N[C@H]2N(C1=O)c1ccccc1[C@]21C[C@@H]2C(=O)N[C@](C)(O1)c1nc3ccccc3c(=O)n12 | -15.12 | 443.15935415200056 | 54 | 1083809824 |
| COc1ccc2c3c1O[C@H]1[C@]4(OC)C=C[C@@]5(c6c4c(O)c4ccccc4c6O)[C@@H](C2)N(C)CC[C@]315 | -9.969 | 469.18892296400077 | 62 | 1500940775 |
| O=C1[C@H]2[C@@H](C(=O)N1c1ccc(F)cc1)[C@@H]1CCCN1[C@@]21c2ccccc2-n2c1nc1ccccc1c2=O | -9.419 | 492.1597687520006 | 58 | 35385143 |
| O=C1c2ccccc2[C@]2(O)Nc3nc4c(ncn4[C@@H]4O[C@H](CO)[C@H](O)[C@H]4O)c(=O)n3[C@@]12O | -9.321 | 443.10771250400035 | 49 | 247857373 |
| CC1(C)C(=O)N2c3ccccc3[C@]3(C[C@@H](n4cnc5ccccc5c4=O)C(=O)O3)[C@H]2N1O | -9.312 | 432.1433697400005 | 52 | 85564482 |
| CC1(C)C(=O)N2c3ccccc3[C@]3(C[C@H](n4cnc5ccccc5c4=O)C(=O)O3)[C@H]2N1O | -9.233 | 432.1433697400005 | 52 | 85564469 |
| O=C1[C@H]2[C@@H](C(=O)N1c1ccc(F)cc1)[C@H]1CCCN1[C@@]21c2ccccc2-n2c1nc1ccccc1c2=O | -9.088 | 492.1597687520006 | 58 | 35385140 |
| O=C1c2ccccc2[C@]2(O)Nc3nc4c(ncn4[C@@H]4O[C@H](CO)[C@H](O)[C@@H]4O)c(=O)n3[C@@]12O | -9.019 | 443.10771250400035 | 49 | 247857382 |
| O=C1c2ccccc2[C@]2(O)Nc3nc4c(ncn4[C@H]4O[C@@H](CO)[C@@H](O)[C@H]4O)c(=O)n3[C@@]12O | -9.004 | 443.10771250400035 | 49 | 100487562 |
| Cc1ccc(N2C(=O)[C@H]3[C@@H](C2=O)[C@H]2CCCN2[C@@]32c3ccccc3-n3c2nc2ccccc2c3=O)cc1 | -8.937 | 488.1848406280007 | 61 | 39954578 |
| O=C1c2ccccc2[C@]2(O)Nc3nc4c(ncn4[C@@H]4O[C@H](CO)[C@H](O)[C@@H]4O)c(=O)n3[C@]12O | -8.927 | 443.10771250400035 | 49 | 247857380 |
| N=C(N)Nc1ccc2c3c([nH]c2c1)[C@@H]1Oc2c(O)ccc4c2[C@@]12CCN(CC1CC1)[C@H](C4)[C@]2(O)C3 | -8.818 | 471.2270397880008 | 64 | 13863070 |
| O=C1c2ccccc2[C@@]2(O)Nc3nc4c(ncn4[C@H]4O[C@@H](CO)[C@@H](O)[C@H]4O)c(=O)n3[C@]12O | -8.745 | 443.10771250400035 | 49 | 100487547 |
| COc1ccc2c3c1O[C@@H]1[C@]34CCN(C)[C@H](C2)[C@]42C=C[C@@]1(OC)c1c2c(O)c2ccccc2c1O | -8.717 | 469.1889229640007 | 62 | 252494996 |
| C/C=C1/CN2[C@@H]3C[C@@]45c6ccccc6N(COC)[C@]4(O3)[C@H]2C[C@H]1[C@H]5C(=O)OC | -8.683 | 382.1892573120007 | 54 | 1772820109 |
| C/C=C1/CN2[C@@H]3C[C@@]45c6ccccc6N(COC)[C@]4(O3)[C@H]2C[C@H]1[C@@H]5C(=O)OC | -8.641 | 382.1892573120007 | 54 | 1772820110 |
| COc1ccc2c3c1O[C@H]1[C@@]4(OC)C=C[C@@]5([C@@H](C2)N(C)CC[C@]315)n1c2ccccc2c(=O)n14 | -8.605 | 443.18450628000073 | 58 | 1857795661 |
| C/C=C1/CN2CC[C@]34c5ccccc5N(C)[C@@]35OC[C@@]4(C(=O)OC)[C@H]1C[C@@H]25 | -8.599 | 366.19434269200065 | 53 | 1772820186 |
| C[C@@H]1CC(=O)C2=C3C(=O)c4cccc(O)c4[C@H](O)[C@]3(O)[C@H](O)[C@@H]3OC(=O)[C@@H]4CCCN4[C@@]23C1 | -8.555 | 455.15801676000063 | 58 | 1772591891 |
| O=C1c2ccccc2[C@]2(O)Nc3nc4c(ncn4[C@H]4O[C@@H](CO)[C@@H](O)[C@H]4O)c(=O)n3[C@]12O | -8.552 | 443.10771250400035 | 49 | 100487557 |
| O=C1c2ccccc2[C@@]2(O)Nc3nc4c(ncn4[C@H]4O[C@@H](CO)[C@@H](O)[C@H]4O)c(=O)n3[C@@]12O | -8.483 | 443.10771250400035 | 49 | 100487552 |
| Cc1ccccc1N1C(=O)[C@@H]2[C@@H]3CCCN3[C@]3(c4ccccc4-n4c3nc3ccccc3c4=O)[C@@H]2C1=O | -8.429 | 488.18484062800064 | 61 | 25107422 |
| O=C1c2ccccc2[C@]2(O)Nc3nc4c(ncn4[C@@H]4O[C@H](CO)[C@H](O)[C@H]4O)c(=O)n3[C@]12O | -8.41 | 443.10771250400035 | 49 | 247857377 |
| COc1ccc2c3c1O[C@@H]1[C@@]4(OC)C=C[C@@]5([C@@H](C2)N(C)CC[C@]315)n1c2ccccc2c(=O)n14 | -8.408 | 443.18450628000073 | 58 | 1857795660 |
| COc1ccc2c3c1O[C@@H]1[C@@]4(OC)CC[C@@]5([C@@H](C2)N(C)CC[C@]315)n1c2ccccc2c(=O)n14 | -8.389 | 445.2001563440008 | 60 | 1857795682 |
| COc1ccc2c3c1O[C@H]1[C@@]4(OC)CC[C@@]5([C@@H](C2)N(C)CC[C@]315)n1c2ccccc2c(=O)n14 | -8.367 | 445.2001563440008 | 60 | 1857795683 |
| COc1ccc2c3c1O[C@]1(C)[C@]4(OC)C=C[C@]5(c6c(O)ccc(O)c64)[C@@H](C2)N(C)CC[C@]351 | -8.367 | 433.18892296400077 | 59 | 103845146 |
| COc1ccc2c3c1O[C@H]1[C@]4(OC)C=C[C@@]5(c6c4c(OC)c4ccccc4c6OC)[C@@H](C2)N(C)CC[C@]315 | -8.366 | 497.2202230920009 | 68 | 1500940773 |
| Oc1ccc2c3c1O[C@@H]1c4[nH]c5ccccc5c4C[C@]4(O)[C@@H](C2)N(CC2CC2)CC[C@]314 | -8.358 | 414.19434269200076 | 57 | 148996341 |
| COC(=O)C1=C(N)Oc2[nH]nc(-c3ccccc3)c2[C@@]12C(=O)N(Cc1ccc(F)cc1)c1ccccc12 | -8.333 | 496.1546833720006 | 58 | 9328597 |
| COc1ccc2c3c1O[C@@]1(C)[C@]4(OC)C=C[C@]5(c6c(O)ccc(O)c64)[C@@H](C2)N(C)CC[C@]351 | -8.311 | 433.18892296400077 | 59 | 103845141 |
| N=C(N)Nc1ccc2c3c([nH]c2c1)[C@@H]1Oc2c(O)ccc4c2[C@@]12CCN(CC1CC1)[C@@H](C4)[C@]2(O)C3 | -8.273 | 471.2270397880008 | 64 | 100764695 |
| C[C@@H]1CC(=O)C2=C3C(=O)c4cccc(O)c4[C@H](O)[C@]3(O)[C@H](O)[C@@H]3OC(=O)[C@H]4CCCN4[C@]23C1 | -8.154 | 455.15801676000063 | 58 | 1772591888 |
| O=C1[C@H]2[C@@H](C(=O)N1c1ccccc1F)[C@H]1CCCN1[C@@]21c2ccccc2-n2c1nc1ccccc1c2=O | -8.109 | 492.1597687520006 | 58 | 101500434 |
| C=CCN1C(=O)[C@]2(C(C(=O)c3cc4ccccc4o3)=C(O)C(=O)N2CCN(C)C)c2ccccc21 | -8.099 | 471.1794209000007 | 60 | 19897792 |
| C[C@@H]1CC(=O)C2=C3C(=O)c4cccc(O)c4[C@H](O)[C@]3(O)[C@H](O)[C@@H]3OC(=O)[C@H]4CCCN4[C@@]23C1 | -8.089 | 455.15801676000063 | 58 | 1772591890 |
| COc1ccc(OC)c2c1[C@H]1C(=C(C)C)[C@H]2[C@]2(Cl)[C@@]1(Cl)C(Cl)=C(Cl)S2(=O)=O | -8.058 | 467.95234071200036 | 43 | 257346980 |
| Cc1ccccc1N1C(=O)[C@H]2[C@@H]3CCCN3[C@]3(c4ccccc4-n4c3nc3ccccc3c4=O)[C@@H]2C1=O | -8.046 | 488.18484062800064 | 61 | 103508648 |
| COc1ccc(OC)c2c1[C@H]1C(=C(C)C)[C@H]2[C@@]2(Cl)C(Cl)=C(Cl)S(=O)(=O)[C@]12Cl | -8.039 | 467.9523407120004 | 43 | 104294204 |
| N=C(N)Nc1ccc2[nH]c3c(c2c1)C[C@]1(O)[C@@H]2Cc4ccc(O)c5c4[C@]1(CCN2CC1CC1)[C@@H]3O5 | -8.023 | 471.2270397880008 | 64 | 8036036 |
| COc1ccc(OC)c2c1[C@H]1C(=C(C)C)[C@H]2[C@@]2(Cl)[C@]1(Cl)C(Cl)=C(Cl)S2(=O)=O | -8.019 | 467.95234071200036 | 43 | 101050795 |
| COc1ccc(OC)c2c1[C@@H]1C(=C(C)C)[C@@H]2[C@@]2(Cl)C(Cl)=C(Cl)S(=O)(=O)[C@]12Cl | -8.012 | 467.9523407120004 | 43 | 104294216 |
| CCCN1C(=O)[C@@]2(C(C(=O)c3cc4ccccc4o3)=C(O)C(=O)N2CCN(C)C)c2ccccc21 | -8.009 | 473.19507096400076 | 62 | 20439056 |
| CCN1C(=O)[C@@]2(c3ccccc31)c1c(C)nn(-c3ccccc3)c1N=C1C=CC(Cl)=NN12 | -8.005 | 442.1308869080006 | 51 | 13149240 |
| CCC1(CC)C(=O)N2N(C1=O)[C@@]1(OC)C=C[C@]23[C@@H]2Cc4ccc(OC)c5c4[C@]3(CCN2C)[C@@H]1O5 | -8.001 | 465.2263710920009 | 65 | 1857795678 |
| O=C1[C@@H]2[C@@H]3CCCN3[C@]3(c4ccccc4-n4c3nc3ccccc3c4=O)[C@@H]2C(=O)N1c1ccccc1F | -8.001 | 492.15976875200056 | 58 | 108522899 |
| COc1ccc(OC)c2c1[C@H]1C(=C(C)C)[C@H]2[C@@]2(Cl)[C@@]1(Cl)C(Cl)=C(Cl)S2(=O)=O | -7.989 | 467.95234071200036 | 43 | 216450441 |
| COC(=O)C1=C(N)Oc2cc(C)n(C)c(=O)c2[C@]12C(=O)N(Cc1ccc(Cl)cc1)c1ccccc12 | -7.941 | 491.1247984840006 | 57 | 9189930 |
| COc1ccc2c3c1O[C@H]1[C@]4(OC)C=C[C@@]5(c6c(OC)ccc(OC)c64)[C@@H](C2)N(C)CC[C@]315 | -7.911 | 447.2045730280008 | 62 | 1500940776 |
| CCC1(CC)C(=O)N2N(C1=O)[C@@]1(OC)C=C[C@]23[C@@H]2Cc4ccc(OC)c5c4[C@]3(CCN2C)[C@H]1O5 | -7.905 | 465.2263710920009 | 65 | 1857795679 |
| COC(=O)C1=C(N)Oc2[nH]nc(-c3ccccc3)c2[C@@]12C(=O)N(Cc1cccc(F)c1)c1ccccc12 | -7.899 | 496.1546833720006 | 58 | 9244062 |
| CN1C(=O)[C@]2(C(C(=O)c3cc4ccccc4o3)=C(O)C(=O)N2C[C@@H]2CCCO2)c2ccccc21 | -7.88 | 458.14778642400057 | 56 | 9160887 |
| COC(=O)C1=C(N)Oc2cc(C)n(C)c(=O)c2[C@@]12C(=O)N(C(C)C)c1ccccc12 | -7.865 | 409.16377083600065 | 53 | 9189965 |
| CN1C(=O)[C@@]2(C(C(=O)c3cc4ccccc4o3)=C(O)C(=O)N2C[C@H]2CCCO2)c2ccccc21 | -7.843 | 458.14778642400057 | 56 | 9160888 |
| O=C1[C@H]2[C@@H](C(=O)N1c1ccccc1)[C@@H]1CCCN1[C@@]21c2ccccc2-n2c1nc1ccccc1c2=O | -7.788 | 474.16919056400064 | 58 | 253388502 |
| COC(=O)C1=C(N)Oc2cc(C)n(Cc3cccnc3)c(=O)c2[C@]12C(=O)N(C)c1ccccc12 | -7.749 | 458.1590198040006 | 56 | 9189928 |
| CN1C(=O)[C@@]2(c3cc(F)ccc31)N(C)C[C@H](c1ccco1)[C@@]21SC(=S)N(Cc2ccco2)C1=O | -7.676 | 497.0879263400006 | 54 | 20567717 |
| COc1ccc2c3c1O[C@@H]1[C@]34CCN(C)[C@H](C2)[C@]42C=C[C@@]1(OC)c1c2c(OC)c2ccccc2c1OC | -7.674 | 497.2202230920008 | 68 | 252486268 |
| COC(=O)C1=C(N)Oc2cc(C)[nH]c(=O)c2[C@@]12C(=O)N(Cc1ccc(Cl)cc1)c1ccccc12 | -7.665 | 477.10914842000057 | 54 | 9130600 |
| COc1ccc2c3c1O[C@@]1(C)[C@]34CCN(C)[C@H](C2)[C@]42C=C[C@@]1(OC)c1c(O)ccc(O)c12 | -7.649 | 433.1889229640007 | 59 | 253505153 |
| CC[C@]12C[C@]3(C(=O)OC)O[C@H]1[C@@H](O)CN1CCc4c(n3c3cc(OC)ccc43)[C@H]12 | -7.614 | 398.1841719320007 | 55 | 85982438 |
| CCN1C(=O)[C@]2(C(C(=O)c3ccc4c(c3)OCCO4)=C(O)C(=O)N2C[C@H]2CCCO2)c2ccccc21 | -7.608 | 490.17400117200066 | 62 | 9087530 |
| CN1C(=O)[C@]2(C(C(=O)c3cc4ccccc4o3)=C(O)C(=O)N2C[C@H]2CCCO2)c2ccccc21 | -7.596 | 458.14778642400057 | 56 | 9160890 |
| C=CCN1C(=O)[C@]2(C(C(=O)c3ccc(C)o3)=C(O)C(=O)N2CCN2CCOCC2)c2ccccc21 | -7.566 | 477.1899855840008 | 62 | 20436564 |
| COC(=O)C1=C(N)Oc2cc(C)n(Cc3cccnc3)c(=O)c2[C@@]12C(=O)N(C)c1ccccc12 | -7.549 | 458.1590198040006 | 56 | 9189929 |
| COC(=O)C1=C(N)Oc2cc(C)n(Cc3ccccn3)c(=O)c2[C@]12C(=O)N(C)c1ccccc12 | -7.54 | 458.1590198040006 | 56 | 9708757 |
| Oc1ccc2c3c1O[C@@H]1c4[nH]c5ccccc5c4C[C@@]4(O)[C@@H](C2)N(CC2CC2)CC[C@]314 | -7.525 | 414.19434269200076 | 57 | 3874479 |
| CN1C(=O)[C@@]2(c3cc(F)ccc31)N(C)C[C@@H](c1ccco1)[C@@]21SC(=S)N(Cc2ccco2)C1=O | -7.502 | 497.0879263400006 | 54 | 13591445 |
| CO[C@@]12[C@H](COC(N)=O)C3=C(C(=O)C(C)=C(N)C3=O)N1C[C@H]1[C@@H]2N1C(=O)c1ccc(Cl)cc1 | -7.475 | 472.1149620720006 | 54 | 101254433 |
| CN1C(=O)[C@@]2(C(C(=O)c3cc4ccccc4o3)=C(O)C(=O)N2C[C@@H]2CCCO2)c2ccccc21 | -7.464 | 458.14778642400057 | 56 | 9160885 |
| COC(=O)C1=C(N)Oc2cc(C)n(C)c(=O)c2[C@]12C(=O)N(Cc1cccc(C)c1)c1ccccc12 | -7.45 | 471.1794209000007 | 60 | 9742823 |
| CO[C@@]12[C@H](COC(N)=O)C3=C(C(=O)C(C)=C(N)C3=O)N1C[C@@H]1[C@@H]2N1C(=O)c1ccc(Cl)cc1 | -7.431 | 472.1149620720006 | 54 | 101254430 |
| CCN1C(=O)[C@@]2(C(C(=O)OC)=C(N)Oc3cc(C)n(Cc4ccco4)c(=O)c32)c2ccccc21 | -7.426 | 461.1586854560006 | 57 | 9189935 |
| COC(=O)C1=C(N)Oc2cc(C)n(C)c(=O)c2[C@]12C(=O)N(Cc1ccc(F)cc1)c1ccccc12 | -7.42 | 475.1543490240006 | 57 | 9189940 |
| COC(=O)C1=C(N)Oc2cc(C)n(Cc3ccccc3)c(=O)c2[C@]12C(=O)N(C)c1ccccc12 | -7.41 | 457.16377083600065 | 57 | 9742790 |
| CCn1c(C)cc2c(c1=O)[C@]1(C(=O)N(Cc3ccc(F)cc3)c3ccccc31)C(C(=O)OC)=C(N)O2 | -7.396 | 489.1699990880006 | 60 | 9189946 |
| COC(=O)C1=C(N)Oc2cc(C)n(C)c(=O)c2[C@@]12C(=O)N(Cc1cccc(C)c1)c1ccccc12 | -7.386 | 471.1794209000007 | 60 | 9742817 |
| COc1ccc2c3c1O[C@@H]1[C@]34CCN(C)[C@H](C2)[C@]42C=C[C@@]1(OC)c1c(OC)ccc(OC)c12 | -7.377 | 447.20457302800077 | 62 | 252495826 |
| COc1ccc2c3c1O[C@]1(C)[C@H](OC(C)=O)[C@@H](O)C=C4[C@@H](C2)N(C)CC[C@]431 | -7.374 | 371.17327290000077 | 52 | 253504759 |
| C=CC(C)(C)[C@@]12C[C@@]3(O)c4nc5ccccc5c(=O)n4[C@H](C)C(=O)N3[C@@H]1N(C(C)=O)c1ccccc12 | -7.371 | 484.2110553760008 | 64 | 3802698 |
| COC(=O)C1=C(N)Oc2cc(C)n(Cc3ccco3)c(=O)c2[C@]12C(=O)N(C)c1ccccc12 | -7.369 | 447.14303539200057 | 54 | 9189932 |
| CCOC(=O)C1=C(N)Oc2cc(C)[nH]c(=O)c2[C@@]12C(=O)N(Cc1ccccc1Cl)c1ccccc12 | -7.36 | 491.1247984840006 | 57 | 9742975 |
| CCN1C(=O)[C@@]2(C(C(=O)c3ccc4c(c3)OCCO4)=C(O)C(=O)N2C[C@H]2CCCO2)c2ccccc21 | -7.349 | 490.17400117200066 | 62 | 9160476 |
| C=CCN1C(=O)[C@@]2(C(C(=O)OC)=C(N)Oc3cc(C)n(Cc4cccnc4)c(=O)c32)c2ccccc21 | -7.343 | 484.174669868007 | 60 | 9189960 |
| CCCN1C(=O)[C@@]2(C(C(=O)c3ccc(C)o3)=C(O)C(=O)N2CCN2CCOCC2)c2ccccc21 | -7.334 | 479.20563564800085 | 64 | 20393231 |
| CCN1C(=O)[C@@]2(C(C(=O)OC)=C(N)Oc3cc(C)n(C[C@@H]4CCCO4)c(=O)c32)c2ccccc21 | -7.301 | 465.18998558400074 | 61 | 9189953 |
| N=C(N)Nc1ccc2[nH]c3c(c2c1)C[C@@]1(O)[C@H]2Cc4ccc(O)c5c4[C@@]1(CCN2CC1CC1)[C@H]3O5 | -7.3 | 471.2270397880008 | 64 | 3941269 |
| COc1ccc2c3c1O[C@]1(C)[C@@H](OC(C)=O)[C@@H](O)C=C4[C@H](C2)N(C)CC[C@@]431 | -7.295 | 371.17327290000077 | 52 | 96316398 |
| COC(=O)C1=C(N)Oc2cc(C)n(Cc3ccco3)c(=O)c2[C@@]12C(=O)N(C)c1ccccc12 | -7.275 | 447.14303539200057 | 54 | 9189933 |
| O=C1[C@H]2[C@@H](C(=O)N1c1ccc(F)cc1)[C@H]1CCCN1[C@]21c2ccccc2-n2c1nc1ccccc1c2=O | -7.27 | 492.1597687520006 | 58 | 13729662 |
| COc1ccc2c3c1O[C@]1(C)[C@@H](OC(C)=O)[C@H](O)C=C4[C@@H](C2)N(C)CC[C@]431 | -7.262 | 371.17327290000077 | 52 | 253504750 |
| C/C=C1/CN2CC[C@@]34c5cc(O)ccc5N(C)[C@]35OC[C@@]4(C(=O)OC)[C@H]1C[C@H]25 | -7.259 | 382.1892573120007 | 54 | 61997653 |
| COc1ccc2c3c1O[C@]1(C)[C@@H](OC(C)=O)[C@@H](O)C=C4[C@@H](C2)N(C)CC[C@]431 | -7.251 | 371.17327290000077 | 52 | 253504755 |
| C=CCN1C(=O)[C@@]2(C(C(=O)c3cc4ccccc4o3)=C(O)C(=O)N2CCN(C)C)c2ccccc21 | -7.25 | 471.1794209000007 | 60 | 19897791 |
| Cc1ccccc1N1C(=O)[C@H]2[C@@H](C1=O)[C@H]1CCCN1[C@]21c2ccccc2-n2c1nc1ccccc1c2=O | -7.234 | 488.1848406280007 | 61 | 8764995 |
| O=C1[C@H]2[C@@H](C(=O)N1c1ccccc1)[C@H]1CCCN1[C@@]21c2ccccc2-n2c1nc1ccccc1c2=O | -7.231 | 474.16919056400064 | 58 | 39956039 |
| Oc1ccc2c3c1O[C@H]1c4[nH]c5ccccc5c4C[C@@]4(O)[C@@H](C2)N(CC2CC2)CC[C@]314 | -7.224 | 414.19434269200076 | 57 | 3802871 |
| O=C1[C@H]2[C@@H](C(=O)N1c1ccccc1F)[C@@H]1CCCN1[C@@]21c2ccccc2-n2c1nc1ccccc1c2=O | -7.223 | 492.1597687520006 | 58 | 35379731 |
| N=CNc1ccc2[nH]c3c(c2c1)C[C@@]1(O)[C@H]2Cc4ccc(O)c5c4[C@@]1(CCN2CC1CC1)[C@H]3O5 | -7.216 | 456.2161407560075 | 62 | 98052512 |
| C[C@@H]1CC(=O)C2=C3C(=O)c4cccc(O)c4[C@H](O)[C@]3(O)[C@H](O)[C@@H]3OC(=O)[C@@H]4CCCN4[C@]23C1 | -7.205 | 455.15801676000063 | 58 | 1772591889 |
| CCOC(=O)C1=C(N)OC2=C(C(=O)CC(C)(C)C2)[C@@]12C(=O)N(Cc1ccc(F)cc1)c1ccccc12 | -7.172 | 490.19040018400074 | 63 | 27023134 |
| C/C=C1/CN2[C@@H]3C[C@@]45c6ccccc6N(COC)[C@]4(O3)[C@@H]2C[C@@H]1[C@H]5C(=O)OC | -7.166 | 382.1892573120007 | 54 | 1875251951 |
| COc1ccc2c3c1O[C@]1(C)[C@]34CCN(C)[C@H](C2)[C@]42C=C[C@@]1(OC)c1c(O)ccc(O)c12 | -7.162 | 433.1889229640007 | 59 | 73351368 |
| CCOC(=O)C1=C(N)Oc2c(c(C)nn2-c2ccccc2)[C@@]12C(=O)N(C(C)=O)c1ccccc12 | -7.161 | 458.1590198040006 | 56 | 39431426 |
| Cc1ccccc1N1C(=O)[C@@H]2[C@H]3CCCN3[C@]3(c4ccccc4-n4c3nc3ccccc3c4=O)[C@@H]2C1=O | -7.159 | 488.18484062800064 | 61 | 103508650 |
| Cc1ccc(N2C(=O)[C@H]3[C@@H](C2=O)[C@@H]2CCCN2[C@@]32c3ccccc3-n3c2nc2ccccc2c3=O)cc1 | -7.159 | 488.1848406280007 | 61 | 253388510 |
| Oc1ccc2c3c1O[C@H]1c4[nH]c5ccccc5c4C[C@]4(O)[C@@H](C2)N(CC2CC2)CC[C@]314 | -7.156 | 414.19434269200076 | 57 | 148996573 |
| Cc1ccc2c(c1)[C@]1(C(=O)N2C)N(C)C[C@@H](c2ccco2)[C@]12SC(=S)N(Cc1ccco1)C2=O | -7.151 | 493.1129982160007 | 57 | 35855628 |
| COC(=O)C1=C(N)Oc2cc(C)n(C)c(=O)c2[C@@]12C(=O)N(Cc1ccc(F)cc1)c1ccccc12 | -7.15 | 475.1543490240006 | 57 | 9189941 |
| CCN1C(=O)[C@]2(c3ccccc31)c1c(C)nn(-c3ccccc3)c1N=C1C=CC(Cl)=NN12 | -7.134 | 442.1308869080006 | 51 | 13149241 |
| O=C1[C@@H]2[C@@H]3CCCN3[C@]3(c4ccccc4-n4c3nc3ccccc3c4=O)[C@H]2C(=O)N1c1ccccc1F | -7.124 | 492.15976875200056 | 58 | 108522903 |
| CCOC(=O)C1=C(N)OC2=C(C(=O)CC(C)(C)C2)[C@]12C(=O)N(Cc1ccc(F)cc1)c1ccccc12 | -7.122 | 490.19040018400074 | 63 | 27023138 |
| Cc1ccccc1N1C(=O)[C@H]2[C@@H](C1=O)[C@H]1CCCN1[C@@]21c2ccccc2-n2c1nc1ccccc1c2=O | -7.103 | 488.1848406280007 | 61 | 39959031 |
| Cc1ccc(N2C(=O)[C@H]3[C@@H](C2=O)[C@H]2CCCN2[C@]32c3ccccc3-n3c2nc2ccccc2c3=O)cc1 | -7.103 | 488.1848406280007 | 61 | 13727747 |
| CCN1C(=O)[C@]2(C(C(=O)OC)=C(N)Oc3cc(C)n(C[C@@H]4CCCO4)c(=O)c32)c2ccccc21 | -7.091 | 465.18998558400074 | 61 | 9189952 |
| COC(=O)C1=C(N)Oc2cc(C)n(C)c(=O)c2[C@]12C(=O)N(CC(C)C)c1ccccc12 | -7.086 | 423.1794209000007 | 56 | 9742829 |
| CCOC(=O)C1=C(N)Oc2cc(C)n(C)c(=O)c2[C@@]12C(=O)N(Cc1cccc(C)c1)c1ccccc12 | -7.075 | 485.19507096400076 | 63 | 9742998 |
| COC(=O)C1=C(N)Oc2cc(C)[nH]c(=O)c2[C@]12C(=O)N(Cc1ccc(Cl)cc1)c1ccccc12 | -7.052 | 477.10914842000057 | 54 | 9742784 |
| CCOC(=O)C1=C(N)Oc2c(c(C)nn2-c2ccccc2)[C@]12C(=O)N(C(C)=O)c1ccccc12 | -7.044 | 458.1590198040006 | 56 | 39431425 |
| CCN1C(=O)[C@]2(C(C(=O)c3ccc4c(c3)OCCO4)=C(O)C(=O)N2C[C@@H]2CCCO2)c2ccccc21 | -7.039 | 490.17400117200066 | 62 | 9160475 |
| Cc1ccccc1N1C(=O)[C@@H]2[C@@H]3CCCN3[C@]3(c4ccccc4-n4c3nc3ccccc3c4=O)[C@H]2C1=O | -7.025 | 488.18484062800064 | 61 | 248083777 |
| CCN1C(=O)[C@@]2(C(C(=O)c3ccc(C)o3)=C(O)C(=O)N2CCN2CCOCC2)c2ccccc21 | -7.018 | 465.1899855840008 | 61 | 20440090 |
| Cc1ccccc1N1C(=O)[C@H]2[C@H]3CCCN3[C@]3(c4ccccc4-n4c3nc3ccccc3c4=O)[C@@H]2C1=O | -7.001 | 488.18484062800064 | 61 | 25107433 |

Table S2 **Identifiers and metrics of the seven molecules obtained for experimental testing and Anle-138b.** The Vina score is in kcal/mol, and the half time of aggregation is normalised to the negative control (1% DMSO only), which correspondingly has a value of 1. The mean and standard deviation of the normalised half times of aggregation for H1 and Anle-138b across 2 independent experiments are shown. Other values are the mean of technical duplicates in the initial screening experiment.

| Molecule | Identifiers | Metrics | | |
|---|---|---|---|---|
| | SMILES | ZINC id | Vina score | Normalised half time |
| H1 | C[C@@H]1CC(=O)C2=C3C(=O)c4cccc(O)c4[C@H](O)[C@]3(O)[C@H](O)[C@@H]3OC(=O)[C@H]4CCCN4[C@@]23C1 | 1772591890 | -8.717 | 1.6 Â± 0.2 |
| H2 | C[C@@H]1N[C@H]2N(C1=O)c1ccccc1[C@]21C[C@@H]2C(=O)N[C@](C)(O1)c1nc3ccccc3c(=O)n12 | 1083809824 | -15.76 | 0.97 |
| H3 | CCC1(CC)C(=O)N2N(C1=O)[C@@]1(OC)C=C[C@]23[C@@H]2Cc4ccc(OC)c5c4[C@]3(CCN2C)[C@H]1O5 | 1857795679 | -8.156 | 1.01 |
| H4 | COc1ccc2c3c1O[C@H]1[C@@]4(OC)C=C[C@@]5([C@@H](C2)N(C)CC[C@]315)n1c2ccccc2c(=O)n14 | 1857795661 | -8.451 | 1.10 |
| H5 | COc1ccc2c3c1O[C@@H]1[C@@]4(OC)CC[C@@]5([C@@H](C2)N(C)CC[C@]315)n1c2ccccc2c(=O)n14 | 1857795682 | -8.398 | 1.05 |
| H6 | C/C=C1/CN2[C@@H]3C[C@@]45c6ccccc6N(COC)[C@]4(O3)[C@H]2C[C@H]1[C@@H]5C(=O)OC | 1772820110 | -8.672 | 1.06 |
| H7 | O=C1c2ccccc2[C@]2(O)Nc3nc4c(ncn4[C@@H]4O[C@H](CO)[C@H](O)[C@H]4O)c(=O)n3[C@@]12O | 247857373 | -9.333 | 1.06 |
| Anle-138b | Brc1cccc(-c2cc(-c3ccc4c(c3)OCO4)n[nH]2)c1 | 68200503 | n/a | 1.3 Â± 0.1 |
